# A computational model of chemically and mechanically induced platelet plug formation

**DOI:** 10.1101/2023.01.26.525741

**Authors:** Giulia Cardillo, Abdul I. Barakat

**Affiliations:** Hydrodynamics Laboratory, École Polytechnique, Bd des Maréchaux, Palaiseau, 91120, France

**Keywords:** Thrombus formation, shear rate, shear gradient, stenosis

## Abstract

**Objectives:** Thrombotic deposition is a major consideration in the development of implantable cardiovascular devices. Recently, it has been experimentally demonstrated that localized changes in the blood shear rate -i.e. shear gradients-play a critical role in thrombogenesis. The goal of the present work is to develop a predictive computational model of platelet plug formation that can be used to assess the thrombotic burden of cardiovascular devices, introducing for the first time the role of shear gradients. We have developed a comprehensive model of platelet-mediated thrombogenesis which includes platelet transport in the blood flow, platelet activation and aggregation induced by both biochemical and mechanical factors, kinetics and mechanics of platelet adhesion, and changes in the local fluid dynamics due to the thrombus growth.

**Methods:** A 2D computational model was developed using the multi-physics finite element solver COMSOL 5.6. The model can be described by a coupled set of convection-diffusion-reaction equations. Platelet adhesion at the surface was modeled via flux boundary conditions. Using a moving mesh for the surface, thrombus growth and consequent alterations in blood flow were modeled. In the case of a stenosis, the notions of shear stress induced platelet activation in the contraction zone and shear gradients induced platelet deposition in the expansion zone downstream of the stenosis were studied.

**Results:** The model provides the spatial and temporal evolution of platelet plug in the flow field. The computed platelet plug size evolution was validated against literature data. The results confirm the importance of considering both mechanical and chemical aggregation of platelets.

**Conclusions:** The developed model represents a potentially useful tool for the optimization of the design of the cardiovascular device flow path.

## 1 Introduction

Cardiovascular diseases (CVDs) define a group of disorders of the heart and blood vessels that constitute the leading cause of death worldwide [1]. Among CVDs, thrombosis has a significant role, it refers to the formation of a blood clot, known as thrombus, inside a blood vessel, altering or in the worst scenario obstructing the flow through the circulatory system.

The clinical success of many cardiovascular devices is impacted by the thrombotic response.

Most implantable cardiovascular devices, such as prosthetic heart valves, endovascular stents and ventricular-assist devices, inducing hemodynamic alterations, have been shown to promote thrombotic deposition [2],that can lead to the failure of the device. To this regard, thrombosis and thromboembolism remain significant sources of morbidity and mortality in cardiovascular device patients [2].

On the other hand, some devices, such as foam-based or coil-based aneurysm treatments, are designed to promote a controlled thrombus formation process, which leads to the success of the device [3].

In this framework, the availability of a mathematical model capable of reliably predicting regions of platelet activation and deposition in case of hemodynamic alterations, such as within a cardiovascular device, would constitute a powerful tool for optimizing the design of the device itself. This work describes the development of a two dimensional computational model of platelet plug formation which includes some important chemical aspects of the process but also the mechanical ones.

According to Virchow’s triad, there are three possible contributors to excessive and undesirable formation of clots (i.e. thrombosis): vessel wall injury or inflammation, changes in the intrinsic properties of blood, and changes in blood flow field [4].

Thrombus formation –i.e. hemostasis process– is the normal physiological response to vascular injury that prevents significant blood loss [5]. There are two main components of hemostasis: primary hemostasis and secondary hemostasis. Primary hemostasis refers to platelet aggregation and platelet plug formation. Secondary hemostasis refers to the deposition of insoluble fibrin, which is generated by the coagulation cascade. These two processes occur simultaneously and are mechanistically intertwined. Thrombosis can occur whenever these processes are dis-regulated [6].

Platelets are the real protagonists of primary hemostasis. It initiates immediately after vessel injury and the main consequence of this process is the creation of a platelet plug. Primary hemostasis consists of two crucial steps: platelet activation and platelet aggregation. Figure 1 provides a chart that summarizes the processes that lead to platelet plug formation.

**Fig. 1.**
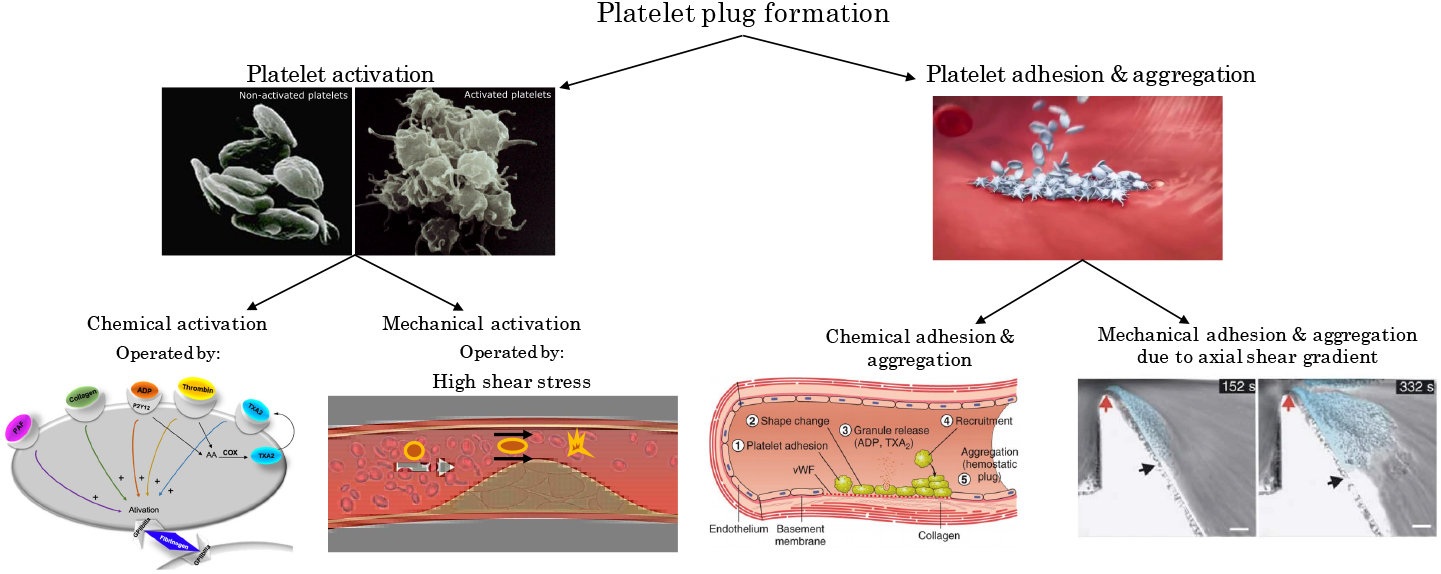
A schematization of the processes involved in platelet plug formation (Figure created using images from [10], [11],[12], [13])

When the endothelial surface of a blood vessel is injured, platelets come into contact with exposed collagen and von Willebrand factor and immediately attach to the injured surface. When seen in fresh blood, platelets appear spheroid, but, in the case of vessel injuries, they have a tendency to extrude hairlike filaments from their membranes – pseudopods– assuming a star-like appearance. This phenomenon is known as platelet activation, see Figure 1. Activated platelets attach to the damaged surface but also to each other in large numbers, forming a tenaciously adherent mass of platelets. They synthesize and release in tiny granules, the agonists, clot-promoting substances that enable the recruitment of other platelets, stimulating the platelet activation itself, namely chemical platelet activation. On the other hand, it has been shown [7],[8],[9] that platelets can also be activated by large mechanical stresses, in particular by high shear stress. In humans, physiologic mean shear stress levels in the arterial circuit reach 20 to 30 dyne/*cm*^2^ (shear rates of whole blood are in the range of 500 to 750 *s*^−1^). Pathologic levels, which can occur in a stenosed artery or in a cardiovascular device, may reach 350 dyne/*cm*^2^ (shear rate larger than 9000 *s*^−1^), and they are associated with mechanical platelet activation [9].

Platelet aggregation is the process of cell-to-cell adhesion, i.e. during aggregation the platelets adhere to other platelets, creating a plug. Chemical aggregation of platelets is mediated by agonists. As seen, agonists activate platelets by binding to specific receptors on the platelet surface, and activated platelets recruit additional platelets to the growing hemostatic plug [14]. In addition to chemical aggregation, it has been recognized that platelet aggregation can also be mediated by mechanical factors, occurring in regions of blood flow disturbance, namely mechanical platelet aggregation.

Nesbitt et al. [12] have shown that shear deceleration due to an expansion of the blood vessel can induce platelet aggregation. More specifically, they have experimentally demonstrated that localized changes in the shear rate along the blood flow, i.e. axial shear gradients, see Figure 2, expose platelets to rapidly changing hemodynamic conditions, leading to the development of stabilized platelet aggregates. Axial shear gradients occur in all the case in which blood shear rate changes along the flow, for instance, in the case of changes in vessel geometry, such as stenosis, see Figure 2, in presence of an aneurysm or within cardiovascular devices.

**Fig. 2.**
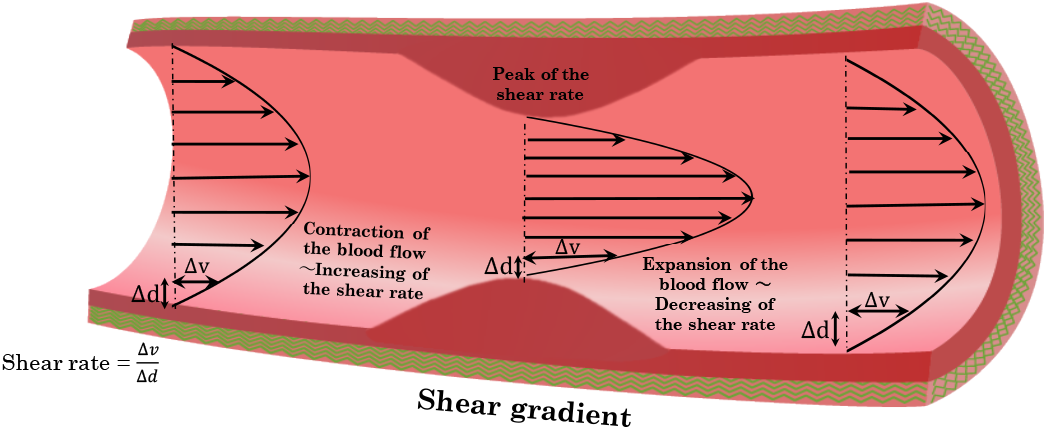
Schematization of axial shear rate changes, i.e.shear gradients, in a stenosed artery

In the past decades, computational modeling has become a very important tool in predicting the formation and the evolution of a thrombus in the cardiovascular system. One of the first continuum models of platelet aggregation was proposed by Fogelson [15],[16]. In this model, resting platelets, activated platelets and one agonist (ADP) are included, and they are regarded as species suspended in the blood whose concentration is obtained by solving the Navier-Stokes equations in combination with the convection-diffusion-reaction equation. Based on this work, Wang and Fogelson [17] developed numerical models, involving a hybrid finite-difference and spectral method, in order to investigate the influence of new chemically-induced activation, link formation, and shear-induced link breaking in determining when aggregates develop sufficient strength to remain intact and when they are broken apart by fluid stresses. Sorensen et al. [2],[18] used Fogelson’s equations as a basis for modeling the concentration of activated and resting platelets as well as ADP [15]; they developed a more complete computational model that simulated platelet-platelet and platelet-surface adhesion, platelet activation by relevant agonists (ADP and TxA2), platelet-phospholipid-dependent thrombin generation, and thrombin inhibition by antithrombin III, ATIII. The latter work is valuable, but it was mainly focused on the chemical aspect of thrombosis. Goodman et al. [19] started from the Sorensen model and added two important fluid dynamical aspects: the influence of thrombus growth on the blood flow and the possibility of embolization due to high shear stress. Hosseinzadegan and Tafti [20] basing their work on Sorensen’s model, proposed a platelets adhesion rate as a linear function of the shear rate. Finally, Wu et al. [21] added to the Sorensen model three important features: the study of thrombus growth, shear-induced platelet activation, and thrombus embolization due to shear.

Wootton et al. [22] drew attention to the effective diffusivity and modeled platelet activation and accumulation at the vessel wall whilst taking into account flow convection and shear-dependent diffusion. This blood clotting model aimed at predicting the formation of a thrombus in a vessel subject to plaque rupture and a lesion in the endothelium. The model of Wootton et al. was further employed by Jordan et al. [23] to investigate the effect of RBC-induced platelet margination and enhanced platelet diffusivity on the adhesion of platelets.

With regard to the influence of thrombus growth on the flow field, a detailed study was proposed by Leiderman and Fogelson [24], who developed a spatio–temporal mathematical model of platelet aggregation and blood coagulation under flow that includes detailed descriptions of coagulation biochemistry, chemical activation and deposition of blood platelets, as well as the two-way interaction between the fluid dynamics and the growing platelet mass, accounting for the porous nature of the thrombus and studying how advective and diffusive transport to and within the thrombus affects its growth at different stages and spatial locations. A further step in this direction was made by the model of Govindarajan et al. [25], which allows the identification of the distinct patterns characterizing the spatial distributions of thrombin, platelets, and fibrin accumulating within a thrombus.

Besides studies, such as the Sorensen’s one, that attribute platelet plug formation to agonists alone, there are some literature studies that propose models of platelet plug formation based mainly on the mechanical aspects. Yazdani et al. [5] developed a shear-dependent platelet adhesive model based on the Morse potential that is calibrated by existing in vivo and in vitro experimental data, coupled with a tissue-factor/contact pathway coagulation cascade. Massai et al. [26] performed a comprehensive analysis of the local hemodynamics within an image-based model of a 51% stenosed internal carotid artery and focused on the influence of disturbed flow caused by the stenosis on flow-induced platelet activation.

Although the existing literature provides valuable models of platelet aggregation and thrombus formation, a thrombogenesis model which takes into account the bio-chemical aspect of the problem, i.e the role of the agonists generated at sites of vascular injury, but also the significant role of axial shear gradients [12] is still lacking. The latter would be expected to become critical in the case of blood flow disturbances, such as in aneuryms, stenosed arteries/arterioles and within cardiovascular devices. To this regard, the present work aims to study platelet aggregate size and platelet deposition at the blood vessel wall, evaluating in detail the role of shear gradients in thrombus formation, including: i) platelet activation, induced by both agonists generated at sites of vascular injury and high shear stress conditions; ii) kinetics and mechanics of platelet-platelet and platelet-surface adhesion; iii) platelet aggregation induced by shear deceleration; iv)alterations in local fluid dynamics due to the presence of a thrombus.

## 2 Methods

The present work is intended to evaluate the platelet deposition at the wall in a blood vessel, introducing and investigating the role of shear gradients in thrombus formation. A two-dimensional model in COMSOL Multiphysics 5.6 was developed.

Schematizing the implementation of the present model, it is composed of four fundamental parts: i) modeling blood flow and species diffusion ii) modeling platelet activation (chemical and mechanical); iii) platelet aggregation (chemical and mechanical); iv) thrombus growth and its influence on the flow field.

The hyphothesis of continuum system is assumed and the platelets along with the agonists and anticoagulant agents were considered as dilute species that are transported by the blood flow.

In particular the model, according to Sorensen et al. work [2],[18], includes seven species: i) resting platelets (*RP*); ii) activated platelets (*AP*); iii) agonist released from platelet granules, such as ADP (*a*_*pr*_); iv) agonist synthesized by the activated platelet, such as thromboxane TxA2 (*a*_*ps*_); v) prothrombin (*PT*); vi) thrombin, generated from prothrombin on platelet phospholipid membranes (*T*); vii) ATIII, antithrombin, which inhibits thrombin and whose action is catalyzed by heparin (*AT*). In order to evaluate the species transport and reactions, for all of them the convection-diffusion-reaction equation is solved:

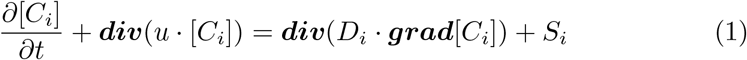

Here, ***div*** is the divergence operator and ***grad*** is the gradient operator. *D*_*i*_ refers to the diffusivity of species i in blood; u is the two-dimensional fluid velocity vector; *C*_*i*_ is the concentration of species i; and *S*_*i*_ is a source term for species i.

Concerning the boundary conditions, for each species a constant physiological concentration was set at the inlet, see in Table 1, while an outflow condition was imposed for all the species at the outlet. At the walls, different kind of flux boundary conditions were considered, as described in section 2.3.

**Table 1.** Physiological values of the main parameters used in the thrombus formation model

| Quantity | Symbol | Value | Unit |
| --- | --- | --- | --- |
| Blood dynamic viscosity [29] | $\mu$ | 0.0035 | Pa · s |
| Blood density [2] | $\rho$ | 1110 | kg/m<sup>3</sup> |
| Platelet Brownian diffusion coefficient [2] | $D_{b,plt}$ | $1.58 \cdot 10^{-9}$ | cm<sup>2</sup> /s |
| ADP Brownian diffusion coefficient [2] | $D_{b,ADP}$ | $2.57 \cdot 10^{-6}$ | cm<sup>2</sup> /s |
| TxA2 Brownian diffusion coefficient [2] | $D_{b,TxA2}$ | $2.14 \cdot 10^{-6}$ | cm<sup>2</sup> /s |
| Thrombin Brownian diffusion coefficient [2] | $D_{b,T}$ | $4.16 \cdot 10^{-7}$ | cm<sup>2</sup> /s |
| Prothrombin Brownian diffusion coefficient [2] | $D_{b,PT}$ | $3.32 \cdot 10^{-7}$ | cm<sup>2</sup> /s |
| Antithrombin Brownian diffusion coefficient [2] | $D_{b,AT}$ | $3.32 \cdot 10^{-7}$ | cm<sup>2</sup> /s |
| Resting Platelet inlet concentration [2] | $[RP]_i$ | $1.5 - 3 \cdot 10^8$ | PLT/ml |
| Activated platelet inlet concentration [2] | $[AP]_i$ | $0 - 0.05[RP]$ | PLT/ml |
| Prothrombin inlet concentration [2] | $[PT]_i$ | 1.1 | μM |
| Antithrombin inlet concentration [2] | $[AT]_i$ | 2.844 | μM |
| Thrombin critical concentration [2] | $[T]_{crit}$ | 0.1 | U ml<sup>-1</sup> |
| TxA2 critical concentration [2] | $[TxA2]_{crit}$ | 0.6 | μM |
| ADP critical concentration [2] | $[ADP]_{crit}$ | 2 | μM |
| Shear rate activation threshold [30],[9] | $\dot{\gamma}_{crit}$ | 10000 | s<sup>-1</sup> |
| Rate of adhesion of resting platelets [18] | $k_{rs}$ | 0.0037 | cm/s |
| Rate of adhesion of activated platelets [19] [18] | $k_{as}$ | 0.135 | cm/s |
| Platelet production rate [31],[32] | $S_p$ | 500 | PLT/(ml · s) |
| Platelet volume [33] | $V_{plt}$ | $4.28 \cdot 10^{-12}$ | cm<sup>3</sup> |

### 2.1 Modeling blood flow and species diffusion

Blood is regarded as an incompressible, Newtonian fluid. The assumption of Newtonian behavior is considered valid at high shear rates 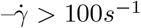 [27]– which are typical of the arterial flows or within cardiovascular devices that are our primary interest. In order to evaluate the blood flow field, the Navier-Stokes equations were solved:

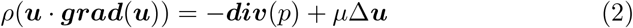

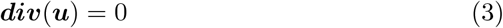

where ***u*** is the flow field, *p* is the pressure and *ρ* and *µ* are the blood density and kinematic viscosity, respectively.

At the inlet, a prescribed constant velocity was imposed, while atmospheric pressure was considered at the outlet. Finally, no slip conditions were imposed at the wall.

Concerning the species diffusion phenomenon, it has been shown that red blood cells have a shear-dependent diffusion-augmenting effect on the transport of platelets and large proteins. To this regard, similarly to Sorensen et al. [2] and Wooton et al.[22], it was decided to adjust the diffusivities *D*_*i*_ of platelets and ”larger” species –i.e.thrombin, prothrombin, and ATIII– adding to the Brownian diffusivity coefficient *D*_*b,i*_, a shear-enhanced diffusivity *D*_*s*_:

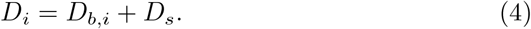

The shear-enhanced diffusivity *D*_*s*_ was modeled considering the Keller formulation [28], in which the augmentation factor is linearly proportional to local shear rate and to red blood cell size:

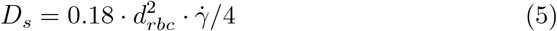

where *d*_*rbc*_ is the diameter of a red blood cell, and 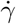 is the local fluid shear rate.

### 2.2 Modeling the platelet activation

Platelet activation was assumed to occur from stimulation by platelet agonists and from shear stress stimulation.

#### 2.2.1 Chemical platelet activation

The chemical platelet activation was simulated following the Sorensen et al. approach, implementing the source terms *S*_*i*_ in eq. 1 for the different species as below:

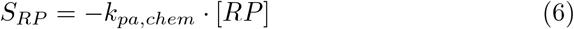

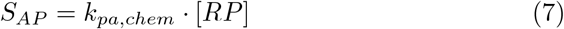

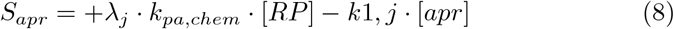

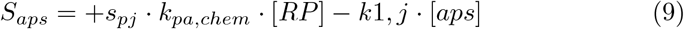

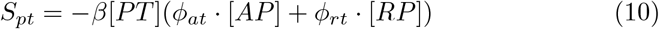

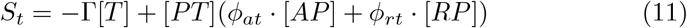

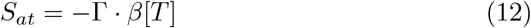

In these equations, *k*_*pa,chem*_ represents a first-order reaction rate constant (in *s*^−1^), which determines the rate of activation of resting platelets–i.e. the rate of consumption of resting platelets and of generation of activated platelets by the first-order reaction *k*_*pa,chem*_[*RP*]– as a function of local agonist concentrations; *λ*_*j*_ is the amount of agonist j released per platelet, so that +*λ*_*j*_ · *k*_*pa*_ · [*RP*] represents the rate at which agonist j is generated from newly activated platelets. The source term for *aps* (Eq.(9)) represents generation and inhibition of an agonist which is synthesized by the activated platelet and the term *s*_*p,j*_ is the rate of synthesis of the agonist. In equation (9) and (8), *k*_1,*j*_ is a first-order reaction rate constant for the inhibition of agonist j.

Basing on Sorensen et al. [2], the rate of activation of resting platelet *k*_*pa,chem*_ was treated as a simplistic linear function of the weighted concentrations of the agonists with an activation threshold:

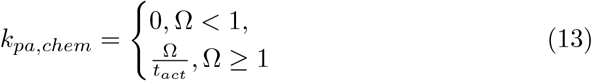

where Ω is the activation function:

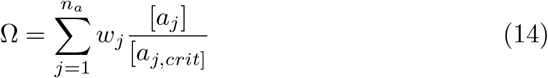

In these equations, *n*_*a*_ is the total number of agonists in the model; *a*_*j*_ is the concentration of the j-th agonist; *a*_*j,crit*_ is the threshold concentration at which the j-th agonist induces platelet activation; *w*_*j*_ is an agonist-specific weight to mimic the differential effects of strong and weak agonist on the activation reaction which here we assume equal to one for all the agonists; and *t*_*act*_ is a “characteristic” time constant for platelet activation in s. In order to avoid function discontinuities, and consequently numerical instabilities, it was decided to model the platelet activation rate as Ω *step*(Ω) where *step*(Ω) is a smooth step function localized where Ω is equal to one and with the minimun and the maximum values equal to zero and one, respectively.

The source terms for PT, T, and AT represent the thrombin-related portion of the model. It is assumed that the generation of thrombin from prothrombin takes place both on platelets adhered at the wall and bulk platelets; the equations for surface-based thrombin generation are incorporated into the flux boundary conditions described in section 2.3. To represent the different rates of thrombin generation from prothrombin at the surface of resting and activated platelets, two rate constants *ϕ*_*rt*_ and *ϕ*_*at*_ in the source terms for PT and T were implemented. In the source terms for T and AT, Γ is the rate of the heparin-catalyzed inactivation of thrombin by antithrombin, which depends on different factors, such as the concentration of heparin [H] (which is assumed constant during the process), the concentration of thrombin and antithrombin (see Appendix A for the expression of Γ).

#### 2.2.2 Mechanical platelet activation

The mechanical platelet activation was implemented through a first-order reaction constant *k*_*pa,mech*_ (in *s*^−1^), which determines a rate of activation of resting platelets as a function of local shear rate. As for *k*_*pa,chem*_ (Eq.(13)), also for modeling *k*_*pa,mech*_ a linear equation with an activation threshold was considered:

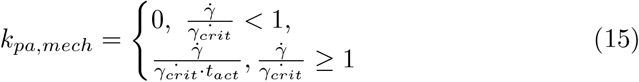

where 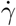 is the shear rate of the blood flow and 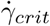 is the critical shear rate at which the resting platelets start to become activated. Thus, in the proposed model, the rate of activation of resting platelets is the sum of the chemical activation rate and the mechanical activation rate:

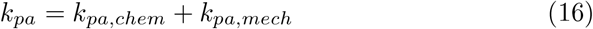

### 2.3 Modeling the platelet adhesion and aggregation

#### 2.3.1 Chemical platelet adhesion and aggregation

The chemical platelet adhesion and aggregation in physiological condition takes place if the endothelium is damaged/injured. The platelets adhering at the surface tend to be activated and they synthesize and release in tiny granules, the agonists that, in turn, make possible the recruitment of other platelets.

Platelet-surface adhesion and platelet-platelet adhesion –i.e. aggregation– are modeled by surface-flux boundary conditions, analogously to Sorensen et al. [2] and Goodman et al. [19].

The flux boundary conditions were described with first-order rate equations as follows (where a positive term indicates an efflux or consumption from the domain and a negative term indicates an influx or generation):

1. adhesion of the resting platelets to surface only, not to other platelets

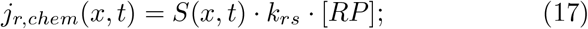
2. adhesion of activated platelets to surface and to deposited activated platelets

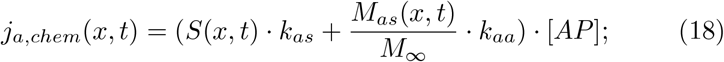
3. generation of platelet-released agonist (*a*_*pr*_)

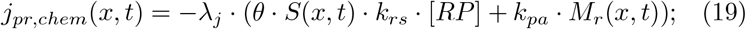
4. generation of platelet-synthesized agonist (*a*_*ps*_)

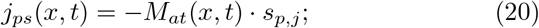
5. consumption of prothrombin due to thrombin generation:

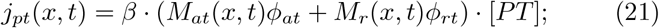
6. generation of thrombin from prothrombin on deposited platelets:

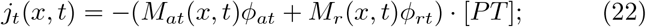

Here, species concentrations are the values at the vessel wall. In these equations, *k*_*rs*_, *k*_*as*_, and *k*_*aa*_ are heterogeneous reaction rate constants that govern the rates of adhesion between, respectively, resting platelets and the surface; activated platelets and the surface; and depositing activated platelets on other platelets; while *θ* is the fraction of adhering resting platelets which activate upon surface contact which can vary between 0 and 1.

All these flux boundary conditions need to be imposed in the portion of the vessel wall that is modeled as injured.

For antithrombin, a no flux condition was imposed.

The platelet-surface deposition equations (17),(18),(19) included a saturation term that limited platelet adhesion to the capacity of the surface, preventing additional platelet-surface adhesion once the capacity of the surface for platelets is reached. The saturation term is defined as follows:

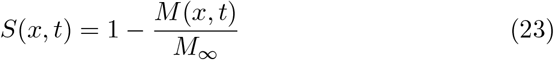

where *M*_*∞*_ is the total capacity of the surface for platelets and *M*(*x, t*) is the total surface coverage with platelets at time *t* and at position *x*, (both in PLT *cm*^−2^) :

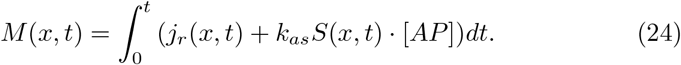

Following Sorensen et al. [2], in addition to *M*(*x, t*), the quantities *M*_*as*_(*x, t*), *M*_*r*_(*x, t*), *M*_*at*_(*x, t*), were defined. They represent respectively the portion of the surface coverage due to activated platelets, the portion of the surface coverage due to resting platelets, the total amount of deposited activated platelets (activated platelet-surface plus platelet-platelet adhesion):

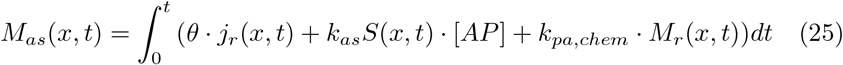

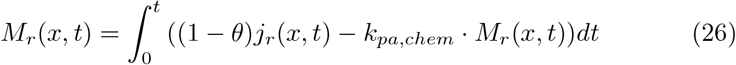

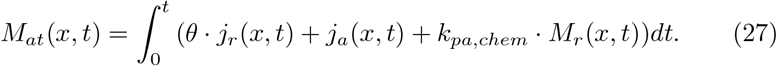

where *k*_*pa,chem*_ is the activation rate constant evaluated at the wall, therefore *k*_*pa,chem*_*M*_*r*_(*x, t*) is the rate of activation of deposited resting platelets due to agonists.

The flux boundary conditions are coupled with the saturation term (Eq.23) and the term *M*(*x, t*), i.e. the total surface coverage with platelets, (Eq.24). Firstly, in order to have a single uncoupled equation that describes the behavior over time of the total surface coverage with platelets, *M*(*x, t*), we combined Equations (17) (23), and (24), obtaining:

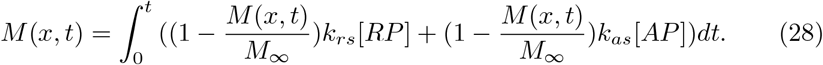

Differentiating Eq.28, it is possible to obtain an ordinary differential equation that describes the evolution of the total surface coverage with platelets over time.

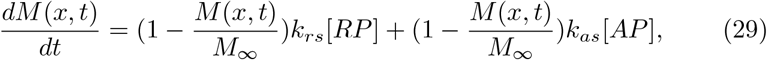

Proceeding analogously for *M*_*as*_(*x, t*), *M*_*r*_(*x, t*) and *M*_*at*_(*x, t*), we obtain:

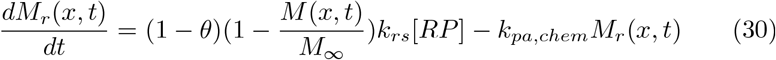

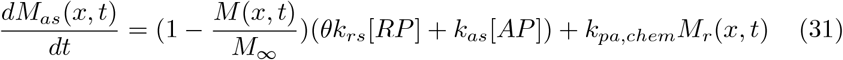

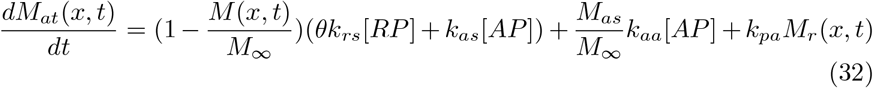

In order to impose the proper flux boundary conditions (Eqs.17-22), the ODE system above has to be solved. Thus, the overall model is represented by a coupled Navier Stokes-CDR-ODE process.

Assuming that *θ* is equal to 1, thus all the adhering resting platelets are activate upon surface contact. The ODE system can be simplified as follows:

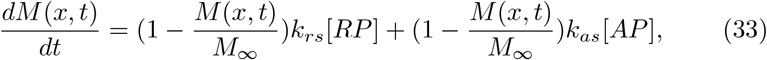

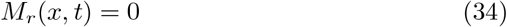

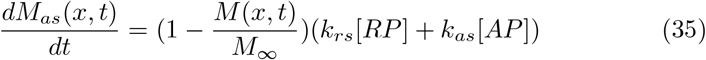

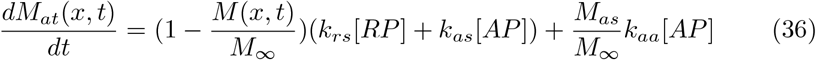

#### 2.3.2 Mechanical platelet adhesion and aggregation

The platelet aggregation due to flow deceleration is here modeled. A flow deceleration translates in a shear rate decrease, i.e. a negative axial derivative of the shear rate. In this perspective, a flux boundary condition for the resting and activated platelets that operates where negative shear gradients occur, is implemented. Furthermore, basing on experimental evidences [12], it appears that platelet aggregation is influenced by the magnitude of the shear gradient. Thus, the introduced flux boundary conditions for the resting and activated platelets are proportional to the absolute value of the negative shear gradient themselves.

The flux boundary condition that mimics the mechanical deposition of resting platelets is:

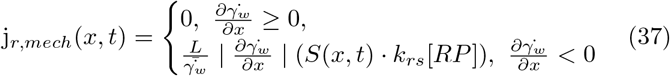

While the flux boundary condition that mimics the mechanical deposition and aggregation of activated platelets is:

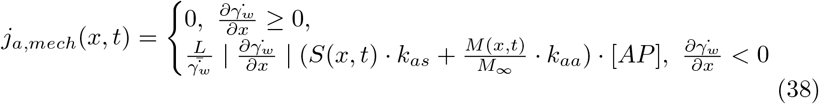

In Eq. 2.3.2-38, the ratio 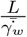 is a scale factor introduced to obtain a more feasible value of the fluxes. Here L is the length of the considered domain and 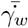 is the wall shear rate in the straight vessel upstream the disturbance.

The flux boundary conditions that characterize the coupled chemical and mechanical problem become:

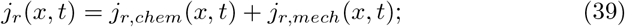

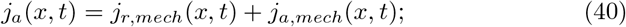

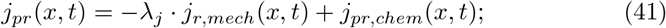

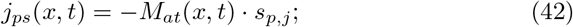

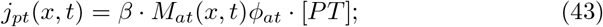

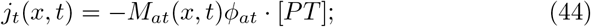

and the ODE system that describes the platelet deposition and aggregation on the damaged surface, including both the chemical and mechanical contributions, becomes:

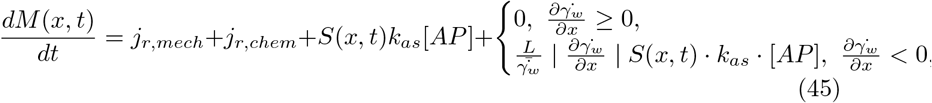

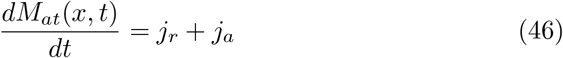

### 2.4 Modeling the influence of thrombus growth on the flow field

In the present model, thrombus growth was tracked with time. An impermeable thrombus was assumed and the associated alterations in local fluid dynamics were determined. To this end, a deformable domain was considered. More specifically, it was decided to parametrize the normal displacement of the domain boundary on which platelets are adhering.

The deformation of the domain, discretized by finite elements, was implemented using the Moving mesh module of COMSOL 5.6 and choosing an Arbirary Lagrangian-Eulerian formulation (ALE).

In the areas of the vessel wall in which the platelets adhere, the normal mesh displacement is imposed as follows:

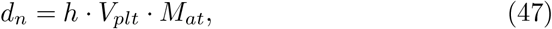

where *V*_*plt*_ is the volume of a single platelet, *M*_*at*_ is the total platelet deposition at the wall and *h* is a scale factor that takes into account the possible presence of red blood cells in the thrombus. With this formulation, at each time step, the total platelet deposition is evaluated and consequently the normal mesh displacement is determined. This mesh displacement influences the flow field at the following time step.

A thrombus can be composed mainly of platelets–termed white thrombus–, or it can be composed of platelets and red blood cells because during its formation and evolution a large number of red blood cells can be trapped.

In order to take into account the presence of red blood cells in establishing the thrombus size, the term *V*_*plt*_ can be increased in Eq. 47. The volume of a single red blood cell is around 24·*V*_*plt*_. Thus, ranging the values of the scale factor *h* from 1 –i.e. a thrombus composed exclusively of platelets– to 24 – i.e. the unrealistic condition of a thrombus composed exclusively by red blood cells– it is possible to model the size of thrombi of different composition.

## 3 Results

The developed computational model allows to evaluate the thrombus formation and evolution associated to chemically and mechanically induced platelet activation and aggregation.

Table 1 provides the physiological values of the main model parameters (see Appendix B for the remaining parameters).

Firstly, the proposed model was validated against literature data. For the model validation, we referred to the *in-vitro* thrombus formation results reported by Nesbitt et al. [12]. In order to reproduce the *in-vitro* conditions, the same wall shear rates measured by the authors in the different sections of their stenosed micro-channel were imposed in the computational model, and basing on the their micro-channel geometry, a domain characterized by a backward-facing step of 90 % stenosis, followed by an expansion zone was designed.

In this simulation, a scale factor *h* of 1 was considered, in accordance with Nesbitt et al. that, in their *in-vitro* experiments observed a thrombus composed mainly of platelets [12].

In Figure 3, the computational predictions in terms of evolution of the aggregate size along the time were compared with the experimental data reported by Nesbitt et al. [12].Thus, the thrombus formation prediction reported in Figure 3 is not the product of a fitting procedure, but the result of the computational model, setting physiological values for the model parameters, see 1. A good agreement between the experimental literature data and the computational results was observed.

**Fig. 3.**
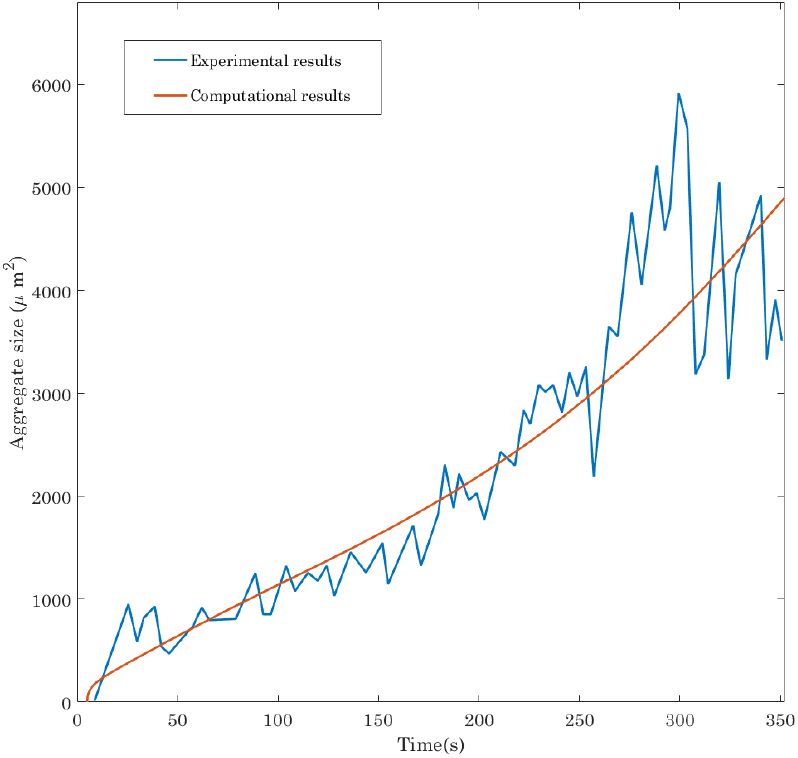
Comparison of the evolution of the experimental aggregate size along the time reported in [12] with computational predictions obtained using the current thrombosis model.

The role of the percentage of stenosis in the thrombus formation was accurately investigated performing a sensitivity analysis that includes an artery characterized by different percentage of stenosis.

The sensitivity analysis was performed basing on the domain reported in Figure 4, panel a. The percentage of stenosis *Ps* is defined as follows:

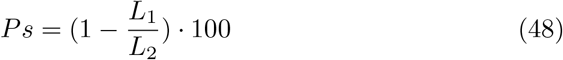

Panel b and c of Figure 4 show the shear rate distribution and the shear gradient distribution in a stenosis, respectively. The contraction demarcates the region of shear acceleration (positive shear gradient). The throat or apex demarcates the stenotic region at which peak shear is experienced. Finally, the expansion demarcates the region of shear deceleration (negative shear gradient).

**Fig. 4.**
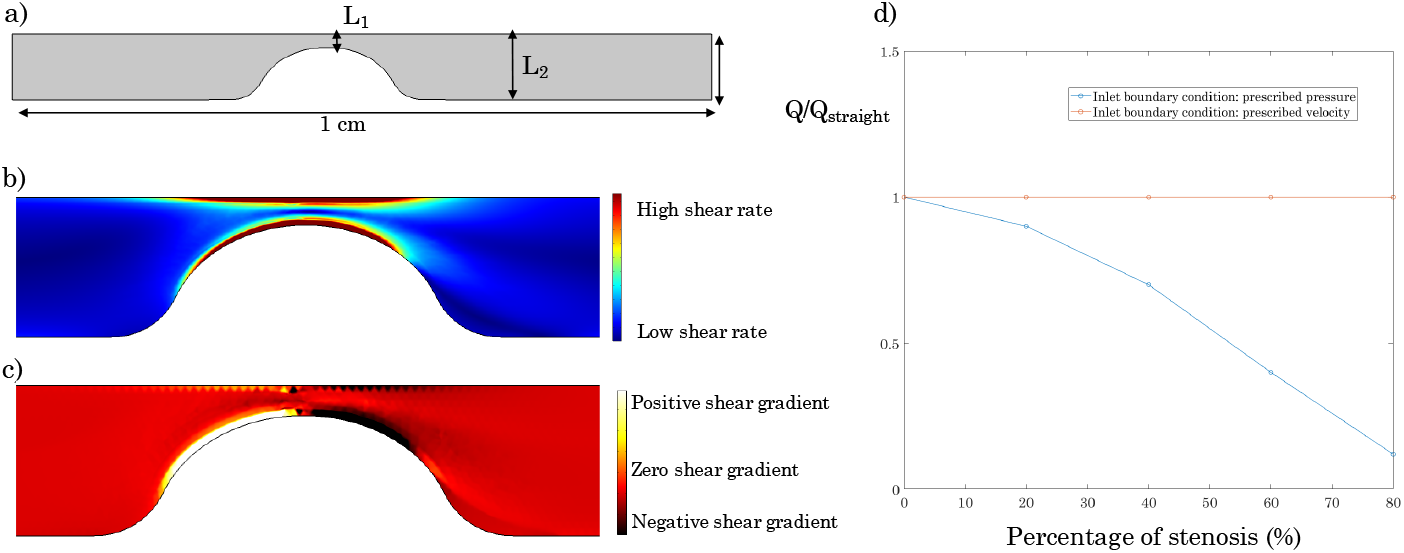
Geometry and inlet boundary conditions considered for the sensitivity analysis of the percentage of stenosis. a)Considered domain, b)Shear rate distribution in a stenosed artery, c) Shear gradients distribution in a stenosed artery, d) Flow rates associated to an inlet prescribed velocity and to an inlet prescribed pressure

Two different inlet boundary conditions were implemented for the blood flow field assessment: a prescribed constant velocity and a prescribed constant pressure.

An inlet prescribed velocity, due to the mass conservation, induces a flow rate that is not influenced by the percentage of stenosis of the artery, Panel d Figure 4. On the other hand, an inlet prescribed pressure induces a flow rate that decreases increasing the percentage of stenosis, as a consequence of the increased hydraulic resistance.

In these simulations, the bottom vessel surface has been modeled as an active surface, i.e. a damaged or inflamed endothelium, and it has been assumed that platelet deposition and aggregation, even if driven by negative shear gradients, can occur only on damaged surface. Furthermore, a scale factor equal to 3 was considered, in order to take into account the platelet plug size’s increase due to the trapping of blood cells during the aggregation process.

An increasing percentage of stenosis, considering an inlet constant velocity (equal to 17 cm/s), induces an increase of both the velocity field and the shear rate in the contraction zone, as reported in Figure 5.

**Fig. 5.**
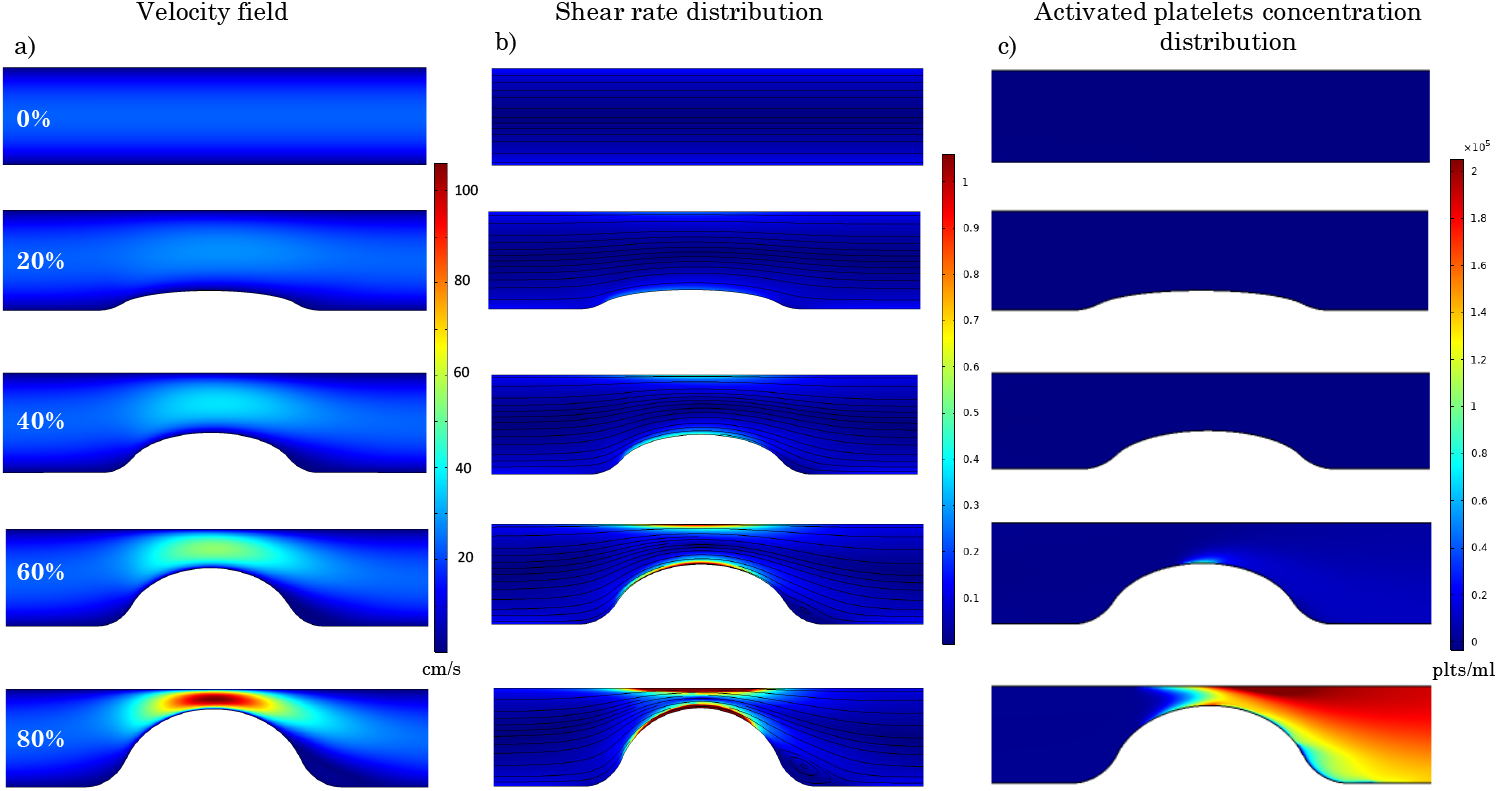
Velocity field, shear rate distribution and platelet activation for different percentage of stenosis associated to a prescribed inlet velocity of 17 cm/s. a)Velocity field for different percentage of stenosis, b) Normalized shear rate distribution and streamlines for different percentage of stenosis, c) Activated platelets concentration distribution for different percentage of stenosis

The streamlines, in panel b of Figure 5, show that recirculation areas are observed in case of a percentage of stenosis of 60 % and 80 %.

The shear rate is normalized basing on the critical shear rate value, implying that values of the shear rate higher than the critical threshold are higher than one.

For values of *Ps* lower than 60 %, the shear rate remains lower than the critical threshold in all the domain.

The critical shear rate (10000 1/s) is reached in the artery characterized by a percentage of stenosis of 60 % and a consequent platelet activation occurs, leading to the activated platelets distribution reported in panel b of Figure 5.

In case of 80% of stenosis, the critical shear rate threshold is reached in a wider area of the domain and the peak of the shear rate encountered is higher than in the 60 % case, resulting in a significantly higher activated platelets concentration, Figure 5.

Imposing a constant inlet velocity condition, a higher percentage of stenosis leads to two effects: first, greater changes in the shear rate along the blood flow direction, i.e. greater magnitude of shear gradients, second, a greater mechanical platelet activation. As a result, an increase of the percentage of stenosis leads to an increase of the platelet plug size, Figure 6.

**Fig. 6.**
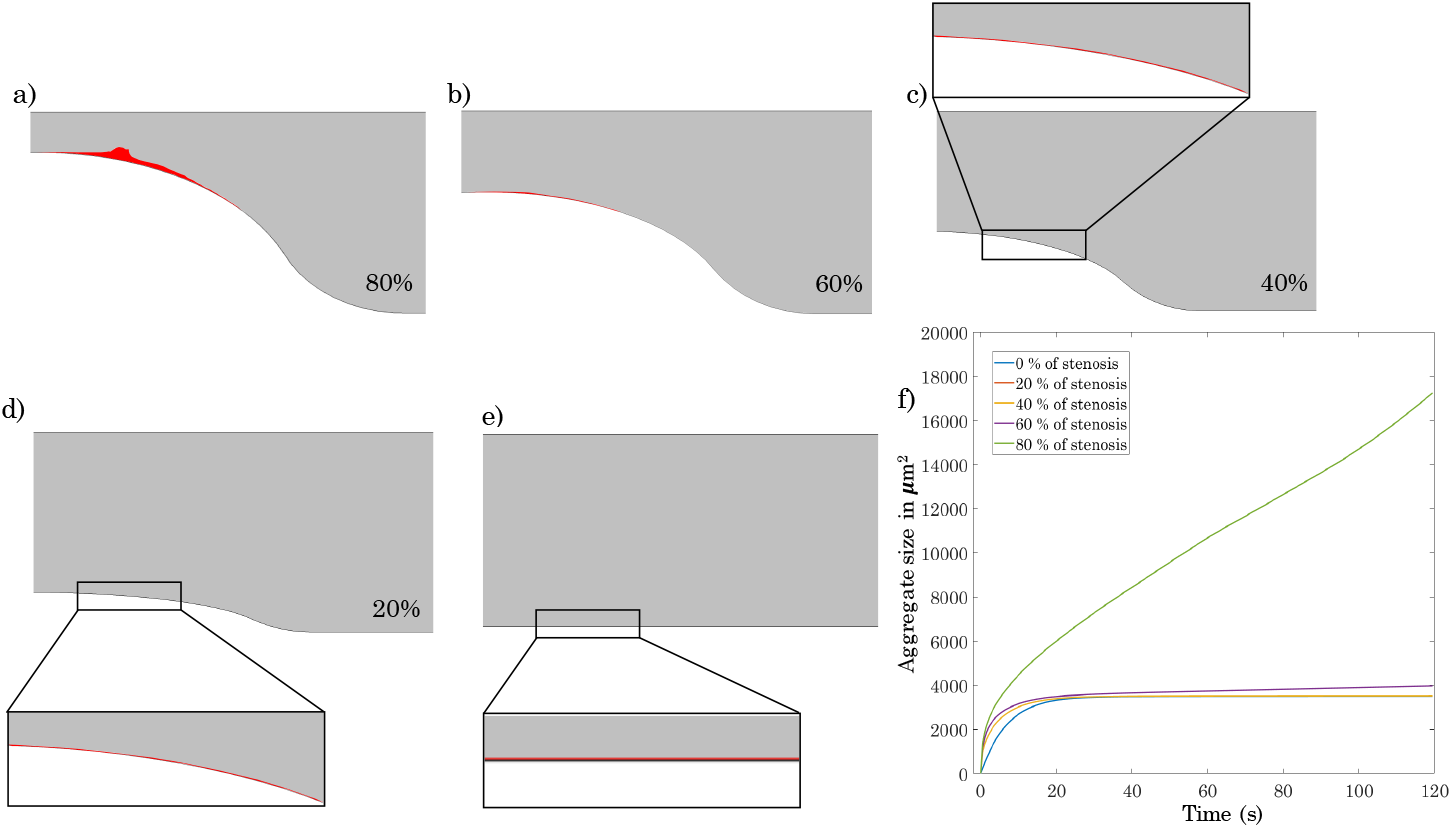
Platelet plug formation for different percentage of stenosis with a prescribed inlet velocity of 17 cm/s and h=3, a),b),c),d),e) platelet plug formation for 80 %, 60%, 40%, 20%, 0% percentage of stenosis, respectively. f) Platelet plug size for different percentage of stenosis along the time

The platelet plug formation associated to an artery characterized by 80 % of stenosis is relevant after only 120 seconds, panel a Figure 6. In this artery configuration, the thrombus evolution along the time (panel f, Figure 6) is greatly faster than the other cases.

The lower platelet activation related to 60 % of stenosis and the lower value of the magnitude of shear gradients induce a smaller platelet plug formation (panel b Figure 6) and the related platelet plug growth rate is much slower (panel f Figure 6) than the previous case. However, the platelet plug size does not reach a plateau, slowly increasing over the simulation time, due to the activated platelets that continue to aggregate to underlying layers of adhered platelets.

Differently, in cases of 40 % and 20 % of stenosis and in case of straight vessel (panel c,d,e of Figure 6,respectively), the platelet plug size reaches a plateau. Indeed, if the shear rate in the contraction zone is not greater than or equal to the critical shear rate, the platelet activation does not occur and once that the damaged surface is saturated with a first layer of resting platelets, the platelet plug growth stops because the resting platelets can not adhere on other platelets.

In Figure 7, a detailed analysis of platelet plug formation in case of 80 % of stenosis is provided. In panel b, it is shown the comparison between the platelet deposition resulted by only biochemical drivers and the platelet deposition resulted by the interplay of biochemical and mechanical drivers. The size of the thrombus promoted only by chemical platelet activation and aggregation is much smaller than the one promoted by chemical and mechanical factors. This comparison seems to confirm the importance of including the mechanical platelet activation and aggregation in a mathematical model, in particular if geometries characterized by non-constant wall shear rate are analyzed.

**Fig. 7.**
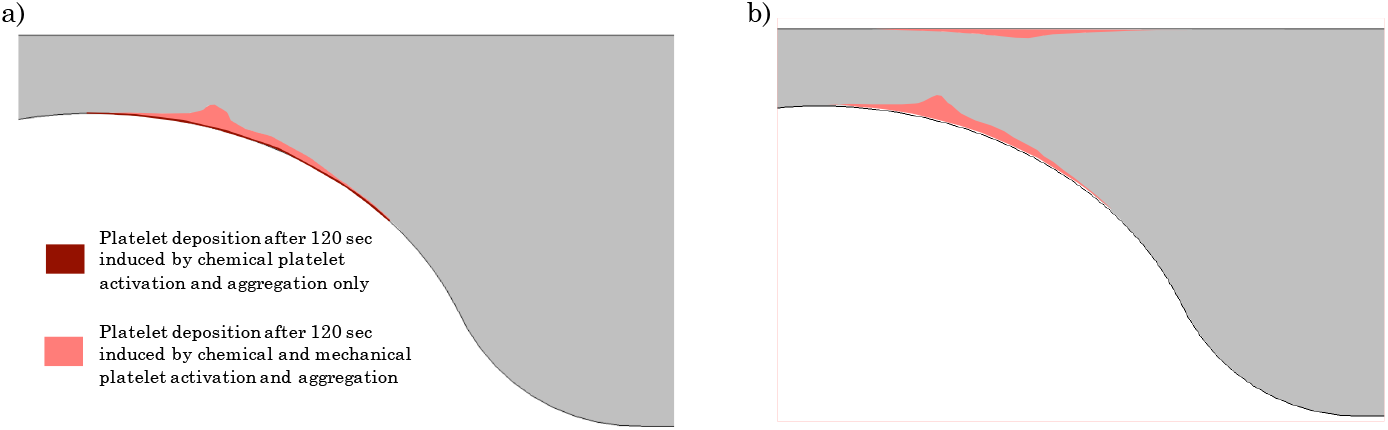
Platelet plug formation in case of 80 % of stenosis after 120 seconds. a) Velocity field in the absence of thrombus formation b) Comparison between the platelet deposition resulted by only biochemical drivers and the platelet deposition resulted by the interplay of biochemical and mechanical drivers. c) Platelet deposition obtained considering that the platelet deposition that

The panel b of Figure 7 reports the thrombus formation that can be observed in case of 80 % of stenosis after 120 seconds, assuming that the platelet aggregation occurs in case of negative shear gradients independently by damaging or inflamming conditions of the endothelium surface. Under this assumption, two areas of platelet deposition are observed, consistently with the negative shear gradients location in case of stenosis, see Figure 4 panel c, as well the *in-vitro* thrombus formation reported by [12].

In Figure 8,the effects of the thrombus growth on the blood flow field in the most critical scenario analyzed are shown. In these conditions, after only 90 seconds the thrombus growth visibly alters the velocity field and after 120 seconds, the velocity field in the artery contraction is significantly changed.

**Fig. 8.**
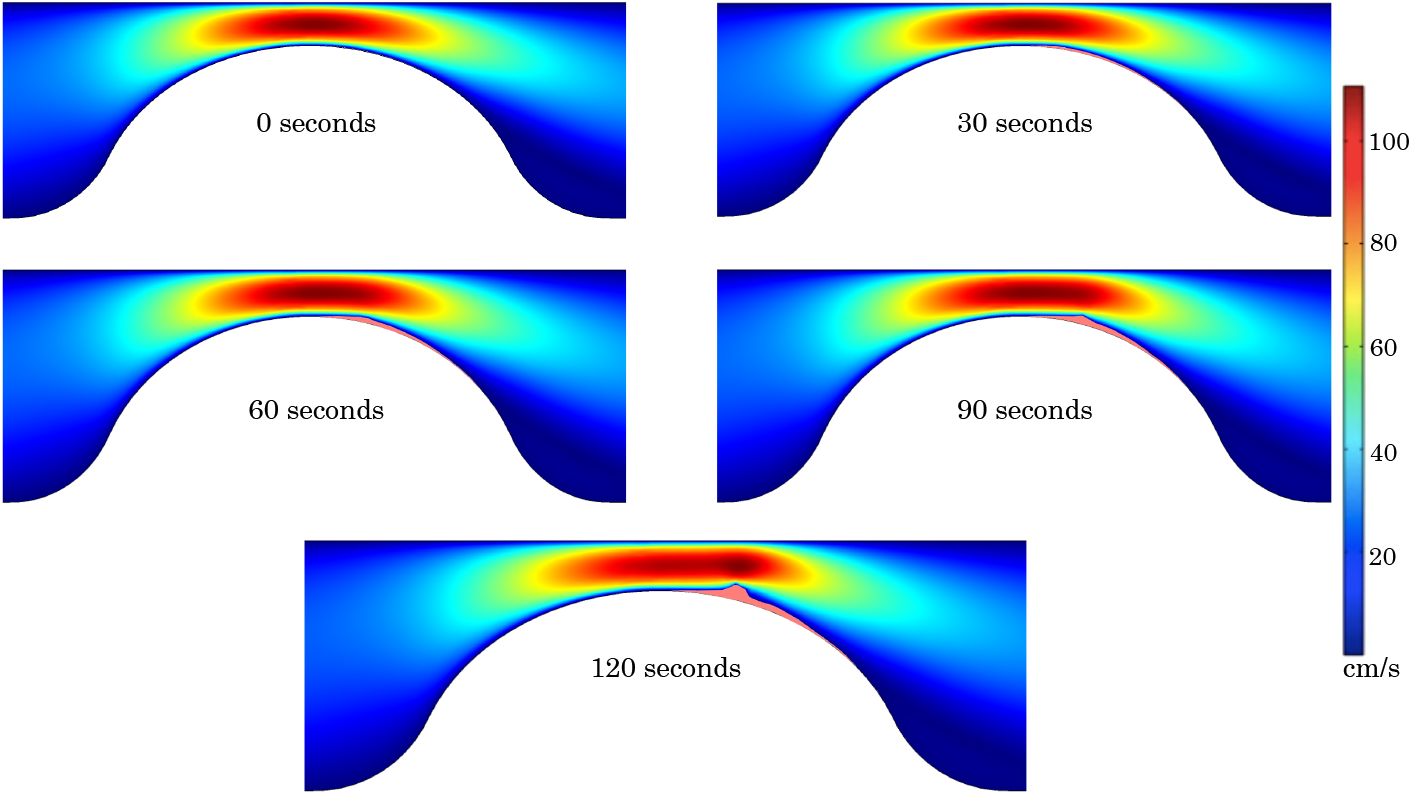
Effect of the thrombus growth on the blood velocity field

Figure 9 reports the blood velocity field, the normalized shear rate with the blood flow streamlines and the platelet activation evaluated in an artery characterized by different percentage of stenosis, considering a constant inlet pressure of 28000 Pa. In an artery with 80% of stenosis, this value of the inlet pressure induces the same blood flow field obtained imposing the inlet velocity of 17 cm/s considered in the previous analysis.

**Fig. 9.**
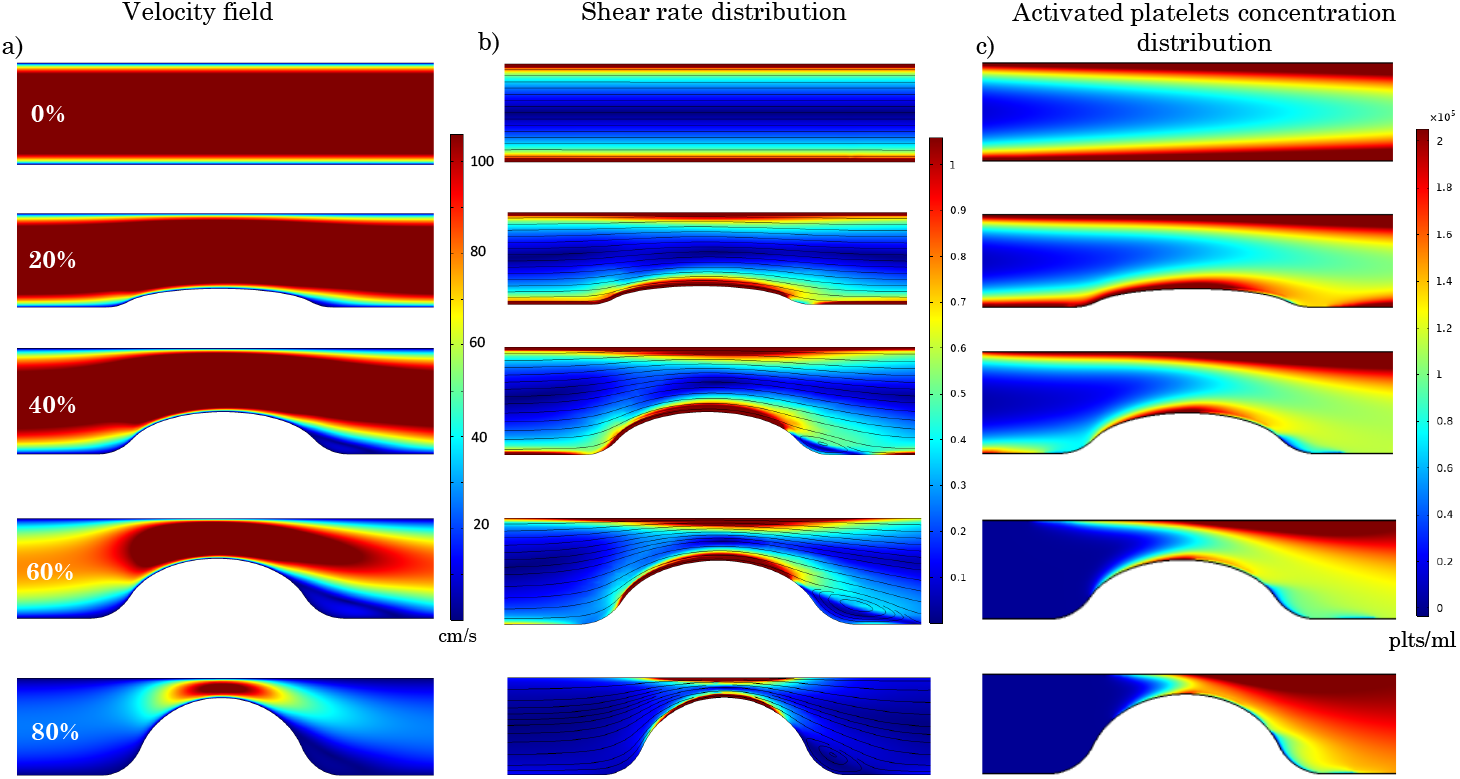
Platelet activation for different percentage of stenosis associated to a prescribed inlet pressure of 28000 Pa. a) Velocity field, b) Shear rate distribution for different percentage of stenosis, c) Activated platelets concentration distribution for different percentage of stenosis

Assuming a constant pressure as inlet boundary condition entails that the wall shear rate decreases increasing the percentage of stenosis. Indeed, the increasing hydraulic resistance linked to more severely stenosed artery, induces a slowdown in the velocity (panel a Figure 9) and a decrease of the flow rate, as reported in panel d of Figure 4.

In these conditions, recirculation areas are observed starting from 40 % of stenosis.

As shown in Figure 9, the considered inlet pressure leads to a wall shear rate value that is higher than the critical one in case of straight vessel. Increasing the percentage of stenosis, the portions of the vessel wall, in which the critical shear rate is reached, are reduced. The platelet activation occurs in any considered artery configurations.

Referring to Figure 10 panel f, it is possible to observe that the aggregate size associated to this prescribed pressure inlet condition increases during the time in any vessel configuration. This behavior is explained by the fact that, as seen in Figure 9, the concentration of activated platelets is important in all the vessel configurations.

**Fig. 10.**
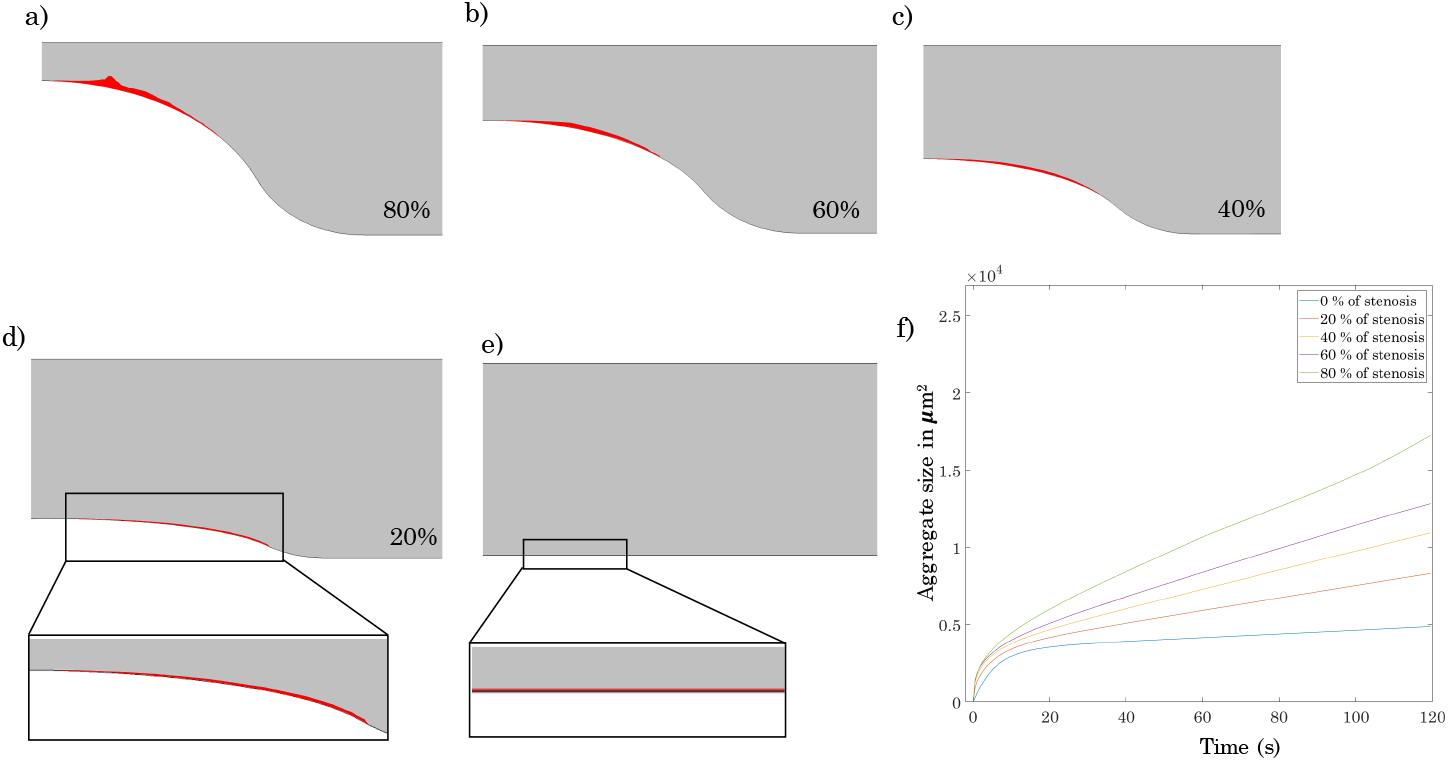
Platelet plug formation after 120 seconds for different percentage of stenosis with a prescribed inlet pressure of 28000 Pa and h=3, a),b),c),d),e) platelet plug formation for 80 %, 60%, 40%, 20%, 0% percentage of stenosis, respectively. f) Platelet plug size for different percentage of stenosis along the time.

The attained aggregate size after 120 seconds in case of 80 % of stenosis is really close that one obtained with a prescribed inlet velocity, having set the boundary conditions in order to have the same initial blood flow field for this case.

Finally, the capabilities of the proposed model were tested in a vessel bifurcation. In particular, the platelet plug response in a bifurcation was investigated for different values of the Reynolds number. The latter analysis assumes that platelet aggregation occurs in case of negative shear gradients, independently by damaging or inflamming conditions of the endothelium surface.

Figure 11 shows the considered bifurcation geometry in panel a. Panel b and c show the shear rate distribution and the shear gradient distribution in the considered bifurcation, respectively.

**Fig. 11.**
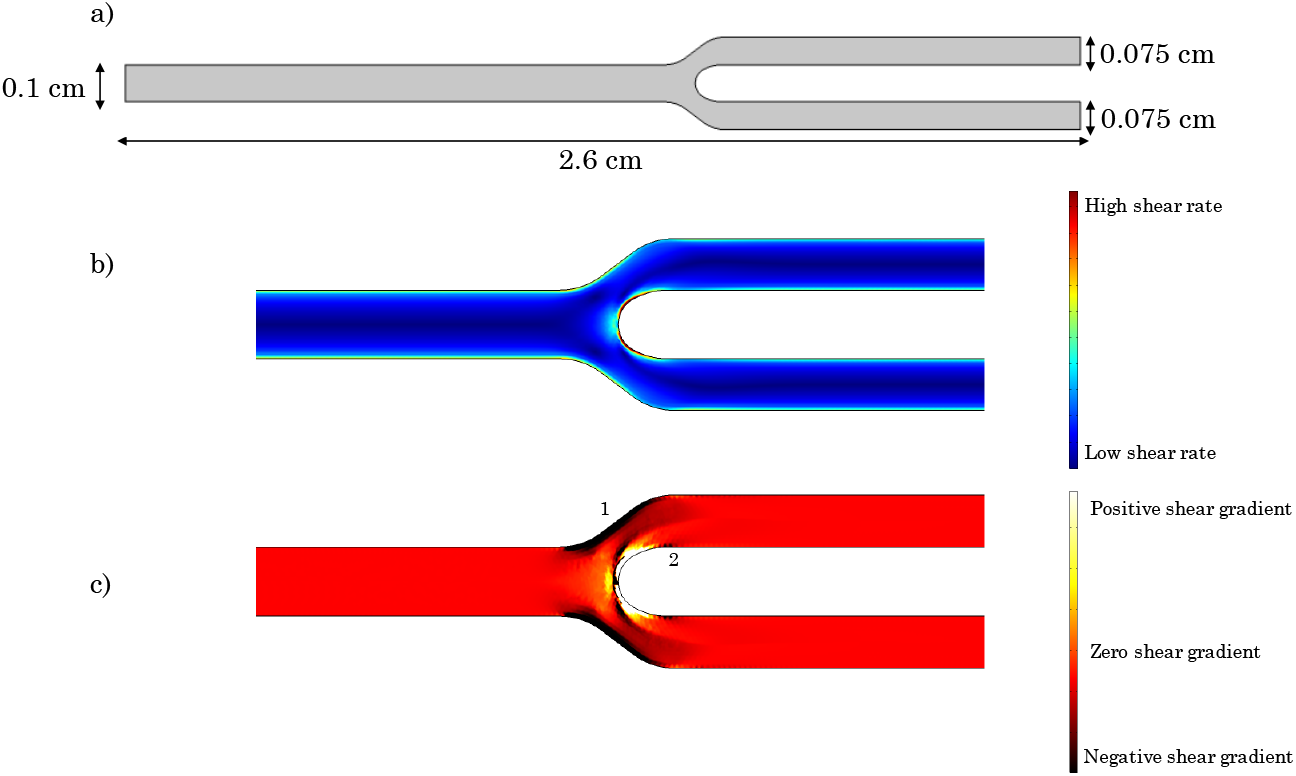
Considered bifurcation geometry and the associated shear rates and shear gradients distributions. a)Domain considered, b)Shear rate distribution in the considered bifurcation, c) Shear gradients distribution in the considered bifurcation.

Due to the well-known skewing of flow toward the inner wall approaching the bifurcation, the highest shear stress zone is present on the inner walls near the branch point, while a lower shear stress zone prevails on the outer walls. Thus, the skewed velocity profile results in the presence of positive shear gradients in the inner wall of the bifurcation and negative shear gradients on outer walls of the bifurcation, zone 1 in panel c Figure 11. Furthermore, after the positive shear gradients area, a flow deceleration is observed, zone 2 of negative shear gradients localization in panel c Figure 11.

The normalized shear rate distribution over the bifurcation and the encountered platelet activation for different values of the Reynolds numbers is reported in Figure 12. The critical shear rate and consequently,the platelet activation don’t occur if the Reynolds numbers is lower than 100. For *Re* equal to 190, platelet activation is localized around the branch point, where the wall shear rate has the peak. Increasing *Re* to 250, the platelet activation is observed in the parent artery walls, around the branch point and downstream in the inner walls of the branches.

**Fig. 12.**
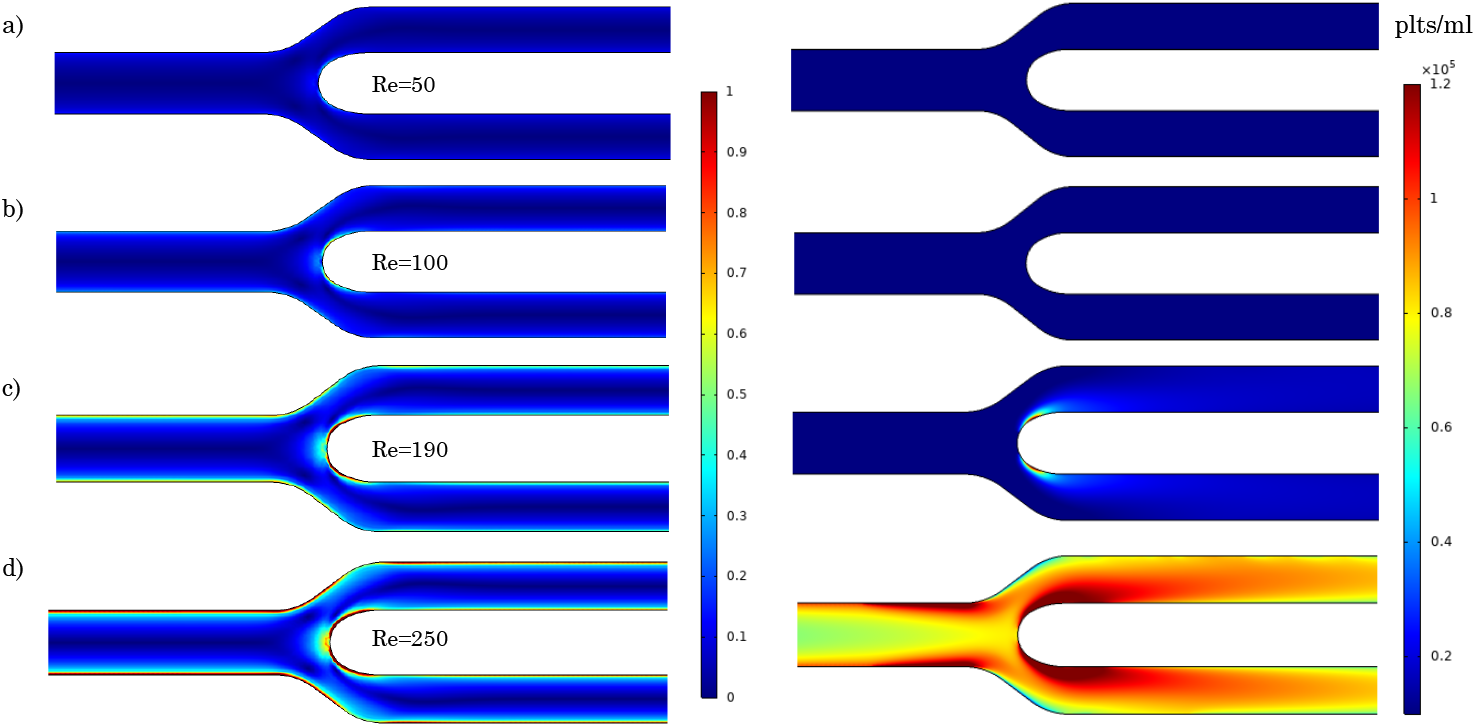
Platelet activation in the considered bifurcation associated to different Reynolds numbers

As regard the platelet plug formation, following the shear gradients distribution, two areas of platelet deposition are observed in the analysed bifurcation, see Figure 13.

**Fig. 13.**
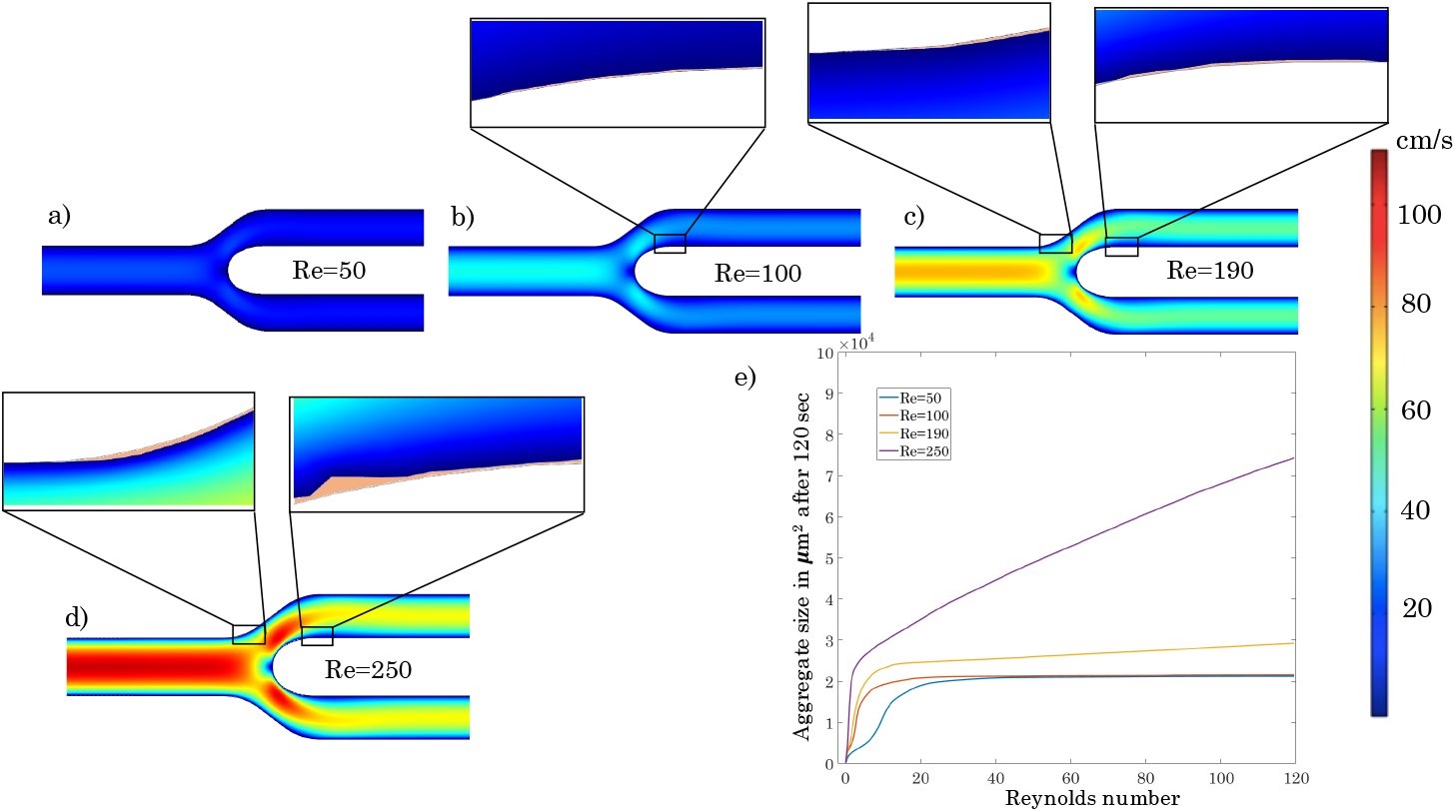
Platelet plug formation in the considered bifurcation for different Reynolds number values after 120 seconds

The platelet plug formation is very small for values of the Reynolds number lower than 100, indeed in these conditions the platelet activation is not reached and once that the portion of the vessel wall characterized by negative shear gradients is saturated with a first layer of resting platelets, the platelet plug growth stops because the resting platelets can not adhere on other platelets. The zoom in Figure 13 shows the platelets monolayer in area 2.

In case of *Re* equal to 190, the plug formation increases downstream the platelet activation areas, i.e. downstream the bifurcation point, in the interior walls of the branches. Whereas in the area 1 (see Figure 11, panel c), the platelet adhesion is limited to a monolayer, because platelet activation occurs only downstream.

Finally, for *Re*=250, after 120 seconds, platelet plug formation starts to be noticeable in correspondence of both zone 1 and 2, in fact in this case platelet activation interests both the parent artery and the branches.

## 4 Discussion and conclusions

The present work provides a computational model capable of predicting platelet plug formation along the vessel wall within a two-dimensional framework, including for the first time the crucial role of the axial shear gradients and evaluating how the platelet aggregate growth itself affects the blood flow field.

According to Virchow’s triad, there are three possible causes of thrombosis onset: vessel wall injury or inflammation, changes in the intrinsic properties of blood, and changes in blood flow field [4]. The proposed model focuses on this latter cause, without disregarding the other two factors of the triad. The model includes the possibility of modeling a vessel wall injury and implements the role of some key biochemical species that are related to thrombus formation.

In physiological conditions, a thrombus is the result of the hemostatic mechanism. In this model, the primary hemostasis that leads to the formation of a platelet plug is implemented. With regard to secondary hemostasis, following the Sorensen et al. [2] approach, it was modeled only partially by introducing the formation of thrombin from prothrombin and its inibhition operated by antitrombin and heparin.

The comparison between the *invitro* platelets plug evolution in time in a stenosed micro-channel [12] and the computationally predicted aggregate size evolution suggests that the model is capable to predict the thrombus formation promoted by the presence of axial shear gradients.

In the context of the model validation, a preliminary analysis was conducted in order to establish the most accurate inlet boundary condition for the convection-diffusion-reaction process. Two type of conditions were analyzed: a constant inlet concentration and a periodic condition.

A constant inlet concentration for the platelets and the other species imply a continuous influx over all the simulation time.

On the other hand, a periodic boundary condition defines a special type of symmetry that obeys cyclic condition across the boundary surface. A periodic condition implies that the same amount of platelets, set as initial condition, passes many times over the domain mimicking the blood circulation.

Figure 14 shows that, even though the platelet plug growth at the beginning is well represented by implementing a periodic condition, after around 50 seconds the aggregate size reaches a plateau because all the platelets in the domain had adhered at the vessel wall.

**Fig. 14.**
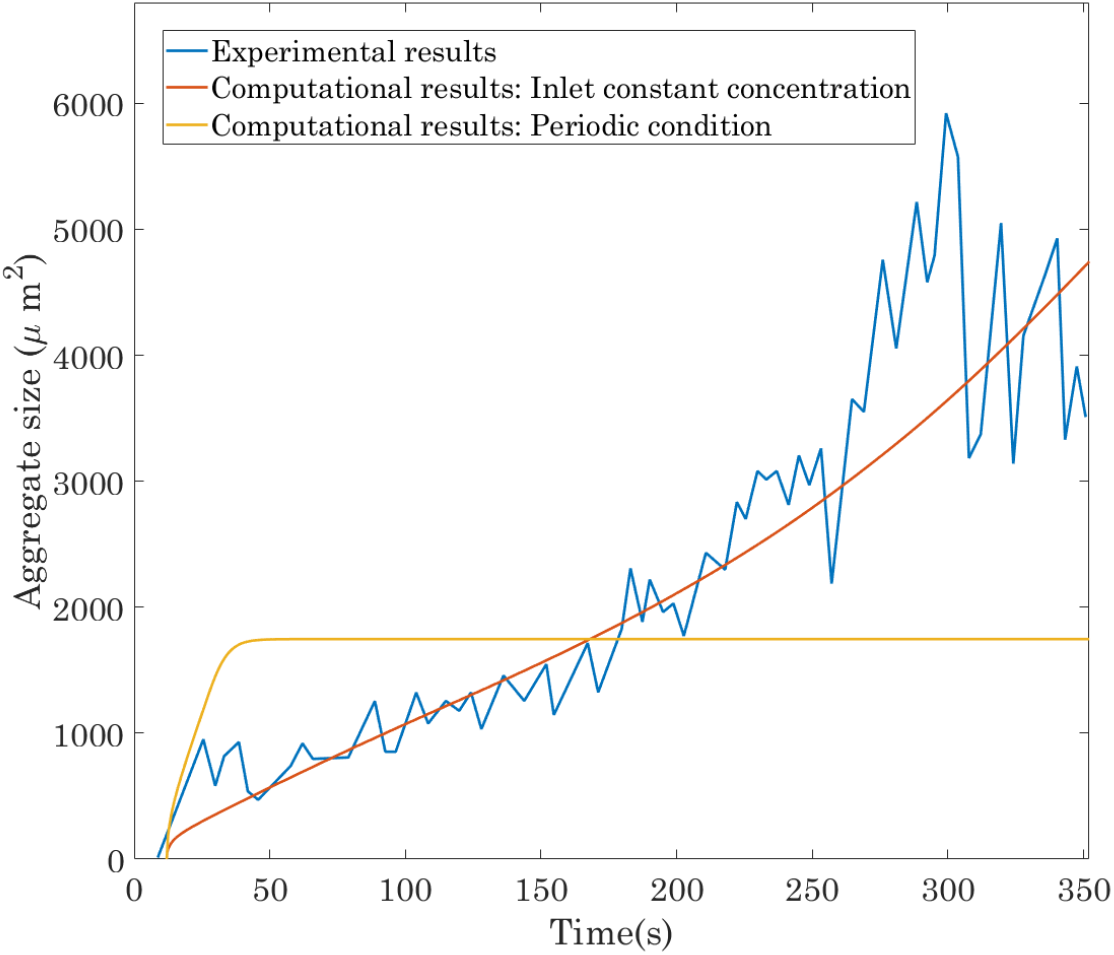
Comparison of the evolution of the experimental aggregate size along the time reported in [12] with computational predictions obtained using the current thrombosis model considering two different inlet conditions for the convection-diffusion-reaction process

This behavior can be explained by the fact that a periodic condition imposes that the same amount of platelets contained in the domain recirculates over all the simulation time. However, if on the one hand it is true that a blood recirculation occur, the total amount of platelets that recirculates is not the only one contained in the domain but also that one contained in the other parts of the system. Thus greater the all system is respect to the analyzed domain, less the periodic condition is able to depict the actual platelet plug growth.

An inlet constant concentration has proven to well describe the platelet plug growth.

As regard the evaluation of the blood flow field, different literature studies have investigated the effects of considering different kind of inlet boundary conditions [34] [35] [36]. In this study two inlet boundary conditions were considered: a constant velocity and a constant pressure.

An inlet constant velocity for the blood flow has been considered in different studies that aim to estimate the thrombus formation [2], [19].

A prescribed velocity at the inlet boundary implies that the blood flow rate does not depend on the percentage of stenosis or, more in general, on the downstream flow disturbances.

An inlet constant pressure implies that increasing the percentage of stenosis and consequently the hydraulic resistance in the domain, the blood flow rate decreases. In the real life, the presence of a severe stenosis may induce a reorganization of the flow field in the upstream arteries and a decrease of the blood flow rate in the stenosed artery, as clinically observed by [37].

In the present study, the two considered inlet flow boundary conditions were set in order to have the same blood flow field in case of 80 % of stenosis. Starting from a percentage of stenosis of 80 %, decreasing the severity of the stenosis leads to a swift decreasing of the platelet plug size considering an inlet velocity. In case of inlet pressure, the platelet plug size reduction is more smooth.

It is known that an injured endothelium attracts a flux of adhering platelets in order to promote the beginning of the hemostatic process [14]. In this computational model, a damaged surface is implemented applying an adhering flux of platelets to the affected vessel walls.

It is not completely understood if in the cardiovascular system, the presence of shear gradients can alone, i.e. without a damaged endothelium, induce thrombus formation [12].

To this regard, for the sensitivity analysis of platelet plug formation to different percentage of stenosis (Figure 6 and Figure 10), the inferior surface were modeled as damaged, assuming that platelet aggregation can occur only on the damaged portion of the vessel wall.

Nonetheless, in order to test the assumption that shear gradients can promote thrombosis independently from the endothelium conditions, we performed two analysis in which no damaged surface were introduced and it was assumed that platelet plug formation can occur in all the areas of the vessel wall interested by negative shear gradients. The first analysis was conducted in 80 % stenosed vessel and the second one in a bifurcation.

In case of 80 % of stenosis and an inlet velocity condition, an analysis without implementing a damaged surface was performed and assuming that platelet plug formation can occur in all the areas of the vessel wall interested by negative shear gradients, Figure 7 panel b. Two areas of platelet plug formation were observed after 120 seconds downstream the stenosis in both lower and upper surface, with a greater aggregate adhering at the lower surface. These results seem in line with the Nesbitt’s in-vitro test [12].

Also the analysis reported in panel a of Figure 7 seems consistent with the Nesbitt’s findings, underlining that if shear micro-gradients occur, then thrombus formation and evolution is driven principally by the mechanical platelet aggregation rather than chemical ones.

Moving to the considered bifurcation geometry, the shear rate distribution reported in Figure 11 agrees with different hemodynamic studies [38],[39].

It is worth pointing out that the platelet deposition, or more in general particles deposition, in a bifurcation are strictly related to the bifurcation configuration, as reported in [40] and [41]. However, the observed location of platelet deposition in the outer walls of the bifurcation (Figure 13) is in line with [42].

It is important to point out that in the analysed bifurcation, the platelet deposition is significant only for Reynolds numbers higher than 100, that are associated to pathological values of the blood flow velocity for the considered geometry.

The implementation of the role of the shear gradients, which was revealed to be fundamental in thrombus formation, is the main novelty of this model. Here, the platelet aggregation induced by shear deceleration is directly proportional to the axial shear-gradient 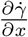, which, resulting from two successive numerical differentiations of the solution of the flow field, can have a noisy output signal. More smooth shear gradient results were obtained by increasing the element order of the fluid dynamics discretization, using P2+P1 elements (P2 for the velocity, and P1 for the pressure). This led to a higher demand on computational resources due to the increase in number of degrees of freedom. The model requires an improvement if precise determination of the shape of the platelet aggregate is desired. Nevertheless, the model has succeeded in predicting the aggregate size in the case of a stenosis. This information may be fundamental in optimizing the design of the flow path of an endovascular device.

A platelet plug formation model that takes into account the chemical and mechanical platelet activation and aggregation and the most important physiological aspects of the problem is computationally very expensive. In this respect, some simplifications were adopted in favor of the numerical tractability, listed below.

First, regarding the geometry, the domains considered are extremely simplified. A step forward would be the implementation of more realistic geometries.

Second, the fibrin formation and its role in the stabilization of the platelet aggregate was not implemented. However, the thrombotic deposition encountered in arteries/arterioles and cardiovascular devices, whose evaluation is the ultimate goal of this model, is characterized by less fibrin involvement than encountered in low-shear venous thrombosis [2].

Thrid, thromboembolism is another feature that was not taken into account in this work. High shear stress has a dual role on thrombus formation and evolution: on the one hand, high shear stresses promote platelet activation, but on the other hand induce the disaggregation of a platelet plug. In the developed model, the thrombus continues to grow until it occludes the blood vessel and the blood flow stops. In the perspective of predicting thrombotic deposition and occlusions, this aspect renders our model very conservative.

Finally, blood is regarded as a Newtonian fluid, this assumption, in case of arterial flow, that is our primary interest, is a good approximation of the real rheological behavior of blood. However, in case of of irregular geometries, such as stenosis, this is a quite strong assumption. It is worth to point out that, in such cases, the shear rate may be locally low, for istance in the recirculation zone. Choi and Barakat [43] investigated the impact of non-Newtonian behavior on the blood flow field over a two-dimensional backward facing step, representing flow disturbance. They found that spatial gradients of shear stress tend to be larger for the non-Newtonian fluid. In this perspective, our model may tend to underestimate the thrombotic deposition.

Despite its simplicity and its limitations, the present work represents an important attempt to couple different key aspects of the thrombus formation and evolution process: chemically and mechanically induced platelet activation and aggregation and the influence of platelet plug growth on the blood flow field, but also on the thrombus evolution itself.

The model promises to serve as a useful tool in helping to predict thrombotic deposition in the cardiovascular system and it may be particularly interesting, from a clinical point of view, to predict thrombus formation in the pathological situations involving blood flow disturbances, such as implanted endovascular devices, aneurysms, stenosis generated by advanced atherosclerotic disease or restenosis due to stent implantation.

The reported model have delivered interesting outcomes and it can lend itself to equally interesting developments. The implementation of more realistic domains that can represent thoroughly the blood flow within and around endovascular devices and the embedding of hemostatic and hemodynamic features that were neglected, are compelling developments of the thrombus formation model. Implementing the shear thinning behavior of blood would allow to estimate more accurately the shear gradient and consequently the thrombotic deposition and to better simulate recirculation zones characterized by locally low shear rate. At the same time, the embedding of the shear thinning rheological behavior of blood as well the role of fibrin stabilization would make the model suitable to evaluate thrombus aggregate size in case of arterial thrombosis but also vein thrombosis. Finally an experimental campaign *ad hoc* would be a beneficial development in order to validate the role of the shear gradients compared to the chemical one and to calibrate more accurately some parameters of the model, such as the scaling term of the thrombus size *h* or the scaling factor of the mechanical aggregation (Eq. (2.3.2)-(38)).

To conclude, the comprehensive thrombus formation model has provided a quantification tool for the thrombotic deposition. To our knowledge, it includes, for the first time, the crucial pro-thrombogenetic role of axial shear gradients, becoming particularly interesting in the pathological scenarios within which blood flow disturbances occur.

## Appendix A Thrombin inhibition operated by Antithrombin

In the source terms for thrombin and antithrombin (Eq.11-12) Γ is the rate of the heparin-catalyzed inactivation of thrombin by antithrombin, which depends on different factors, such as the concentration of heparin [H] (which is assumed constant during the process), the concentration of thrombin and antithrombin

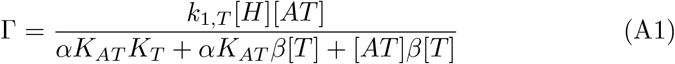

In Eq.A1, *k*_1,*T*_ is a first-order rate constant in *s*^−1^, *K*_*T*_ and *K*_*AT*_ are the dissociation constants (in *µM*) for heparin/thrombin and heparin/ATIII, respectively; *α* represents a factor to simulate a change in affinity of heparin for ATIII when it is bound to thrombin or for thrombin when it is bound to ATIII; and *β* is a conversion factor to convert thrombin concentration from *Uml*^−1^ to *µM*.

## Appendix B Model parameters

The following parameters characterize the part of the model related to chemical platelet activation and aggregation and refer to the Sorensen et al. [2] work.

**Table B1.** Physiological values of the parameters used in the chemical platelet activation and aggregation part of the model.

| Parameter | Value | Unit |
| --- | --- | --- |
| $\lambda_{ADP}$ | $2.4 \cdot 10^{-8}$ | nmol PLT$^{-1}$ |
| $k_{1,TxA2}$ | 0.0161 | s$^{-1}$ |
| $s_{p,TxA2}$ | $9.5 \cdot 10^{-12}$ | nmol PLT$^{-1}$ s$^{-1}$ |
| $k_{1,T}$ | 13.33 | s$^{-1}$ |
| $K_{AT}$ | 0.10 | $\mu M$ |
| $K_T$ | $3.5 \cdot 10^{-2}$ | $\mu M$ |
| $\alpha$ | 1 | [-] |
| $\theta$ | 1 | [-] |
| $\beta$ | $9.11 \cdot 10^{-3}$ | nmol U$^{-1}$ |
| $\phi_{at}$ | $3.69 \cdot 10^{-9}$ | U PLT$^{-1}$s$^{-1}$ ($\mu M$ PT)$^{-1}$ |
| $\phi_{rt}$ | $6.50 \cdot 10^{-10}$ | U PLT$^{-1}$s$^{-1}$ ($\mu M$ PT)$^{-1}$ |
| $t_{act}$ | 1 | s |
| $[H]$ | 2 | $\mu M$ |
| $M_{\infty}$ | $7 \cdot 10^6$ | PLT cm$^{-2}$ |

